# Functional connectivity changes with rapid remission from moderate-to-severe major depressive disorder

**DOI:** 10.1101/672154

**Authors:** Xiaoqian Xiao, Brandon S. Bentzley, Eleanor J. Cole, Claudia Tischler, Katy H. Stimpson, Dalton Duvio, James H. Bishop, Danielle D. DeSouza, Alan Schatzberg, Corey Keller, Keith D. Sudheimer, Nolan R. Williams

## Abstract

Major depressive disorder (MDD) is prevalent and debilitating, and development of improved treatments is limited by insufficient understanding of the neurological changes associated with disease remission. In turn, efforts to elucidate these changes have been challenging due to disease heterogeneity as well as limited effectiveness, delayed onset, and significant off-target effects of treatments. We developed a form of repetitive transcranial magnetic stimulation of the left dorsolateral prefrontal cortex (lDLPFC) that in an open-label study was associated with remission from MDD in 90% of individuals in 1-5 days (Stanford Accelerated Intelligent Neuromodulation Therapy, SAINT). This provides a tool to begin exploring the functional connectivity (FC) changes associated with MDD remission. Resting-state fMRI scans were performed before and after SAINT in 18 participants with moderate-to-severe, treatment-resistant MDD. FC was determined between regions of interest defined a priori by well-described roles in emotion regulation. Following SAINT, FC was significantly decreased between subgenual cingulate cortex (sgACC) and 3 of 4 default mode network (DMN) nodes. Significant reductions in FC were also observed between the following: DLPFC-striatum, DLPFC-amygdala, DMN-amygdala, DMN-striatum, and amygdala-striatum. Greater clinical improvements were correlated with larger decreases in FC between DLPFC-amygdala and DLPFC-insula, as well as smaller decreases in FC between sgACC-DMN. Greater clinical improvements were correlated with lower baseline FC between DMN-DLPFC, DMN-striatum, and DMN-ventrolateral prefrontal cortex. The multiple, significant reductions in FC we observed following SAINT and remission from depression support the hypothesis that MDD is a state of hyper-connectivity within these networks, and rapid decoupling of network nodes may lead to rapid remission from depression.

**Significance statement:** Major depressive disorder is common and debilitating. It has been difficult to study the brain changes associated with recovery from depression, because treatments take weeks-to-months to become effective, and symptoms fail to resolve in many people. We recently developed a type of magnetic brain stimulation called SAINT. SAINT leads to full remission from depression in 90% of people within 5 days. We used SAINT and functional magnetic resonance imaging to determine how the brain changes with rapid remission from depression. We found changes in areas of the brain associated with emotion regulation. This provides a significantly clearer picture of how the non-depressed brain differs from the depressed brain, which can be used to develop rapid and effective treatments for depression.

## Introduction

Major depressive disorder (MDD) is the leading cause of disability worldwide (1, 2) and is increasingly common with a lifetime prevalence in the United States of more than 20% (3). Combating this endemic disease will require improvements in prophylaxis and treatments; however, the development of both is limited in part by insufficient understanding of the neurological changes associated with disease remission.

Investigating the network-level changes associated with remission from MDD has been particularly challenging due to disease heterogeneity of MDD (4, 5), as well as antidepressant medications requiring weeks-to-months to induce remission (6) with limited efficacy (7, 8). Conventional repetitive transcranial magnetic stimulation (rTMS) has been used to investigate functional connectivity changes that occur with remission (9); however, conventional rTMS also requires several weeks to induce remission with less than half of patients remitting (10). Electroconvulsive therapy (ECT) has greater efficacy and more rapid onset of action than antidepressant medications; however, mean onset is still >2 weeks (11), open-label remission rates occur in only 50-65% of patients (7), and there are numerous significant off-target effects, including cognitive disturbances (12) and effects on the motor system (13), making it difficult to determine which network changes are specific to remission from depression. Ketamine induces brief remission rapidly but only in a third of patients (14, 15) and with significant off-target effects (16, 17). Incipient investigations of potent 5-HT_2a_ agonists demonstrate high rates of rapid remission (18, 19); yet, the off-target effects are profound (20). We recently developed a form of repetitive transcranial magnetic stimulation (rTMS) that was associated with remission in 90% of individuals (n=21) with moderate-to-severe, treatment-resistant MDD in a mean of ~3 days without off-target effects in an open-label study (21, 22). This treatment provides a new opportunity to investigate the network alterations associated with remission from MDD.

We chose to begin our investigation at the network level with a resting-state functional connectivity (FC) analysis, because neuroimaging-based approaches that assess resting-state FC have demonstrated some success in investigating neurocircuit-level dysfunction of MDD. These approaches have been used to investigate neurophysiological subtypes of MDD (4, 5) and predict treatment efficacy of rTMS (23, 24, 4, 25–30). Given the promise of this approach, we performed resting state FC analyses on participants before and after SAINT. For this initial investigation, we restricted our analysis to brain regions involved in emotion regulation, as these regions have well-described associations with MDD.

Several lines of evidence suggest dysfunction of emotion regulation in MDD (31) with neural correlates observed with fMRI (32, 33). The neural basis of emotion regulation includes emotional reactivity: amygdala and striatum; explicit control of emotion – central executive network (CEN): dorsolateral prefrontal cortex (dlPFC), ventrolateral PFC (vlPFC); and implicit emotion regulation – default mode network (DMN), salience affective network (SN) and insula. For review, see Etkin et al. (34). Specifically, dysfunction within the DMN, CEN, and SN has repeatedly been reported in individuals with MDD (35–39). Numerous fMRI studies have demonstrated that individuals with MDD have altered frontal connectivity with amygdala (40–44) and striatum (23, 45–47); hyperconnectivity between DMN and amygdala (48–51); and alterations in insular connectivity with multiple nodes (51–53).

Following the emotional dysregulation hypothesis of depression, stimulation of the CEN at the lDLPFC node, via rTMS (37, 54, 55) and transcranial direct current stimulation (56), has been hypothesized to produce antidepressant effects by enhanced regulation of emotion. This is in line with Chen et al’s seminal interleaved TMS/fMRI study demonstrating causal interactions between CEN and DMN, such that increased activity in CEN results in suppression of DMN (57), and this has led others to hypothesize that improved accuracy of targeting these networks might produce better treatment efficacy (27–29, 58–60), as only a third or less of participants remit from their depressive episode with rTMS stimulation of the lDLPFC, a CEN node (61–63). We hypothesized that individualized targeting and enhanced stimulation techniques could improve outcomes (22), and we developed Stanford Accelerated Intelligent Neuromodulation Therapy (SAINT). In our initial open-label trial of SAINT in patients (n=21) with moderate-to-severe, treatment-resistant depression, we observed a strikingly high remission rate of 90% with a mean of ~3 days of treatment to produce remision (21). Thus, FC changes that occur following SAINT may reflect generalized network-level changes associated with remission from depression. Herein, we report our first investigation of the resting-state FC changes that occur in participants with MDD undergoing open-label SAINT.

## Results

### Depression symptoms

Eighteen participants [9 males, mean age 50.8±14.6 (mean±SD)] were recruited for the study and received viable MRI scans before and after SAINT. Participants had a diagnosis of MDD with a mean Hamilton Depression Rating Scale 17-item (HAMD-17) score of 26.3±1.2 and a treatment history of not responding to a mean of 8.7±5.7 antidepressant and 1.3±1.5 augmentation medications. Please refer to the parent clinical trial for more detail on participant characteristics and specific psychotropic medications (21). Participants received a 5-day treatment course under the SAINT protocol. Stimulation location was depth-corrected (64) and chosen based on an algorithm that selected the area within the left dorsolateral prefrontal cortex (lDLPFC) with the most negative FC with sgACC (21). Ten sessions per day of intermittent theta-burst stimulation [iTBS, 60 cycles of 10 bursts of 3 pulses at 50Hz delivered in 2-second trains with an 8-second inter-train interval] were delivered hourly at 90% resting motor threshold for 5 consecutive days for a total of 90,000 pulses.

Mean HAMD-17 significantly decreased from 26.280 (±4.909, range 20-35) at baseline to 4.722 (±4.599, range 0-16) after treatment (t_(17)_=11.275, p<0.001). Mean MADRS significantly decreased from 33.220 (±5.208, range 27-44) at baseline to 4.833 (±5.813, range 0-19) after treatment (t_(17)_=14.915, p<0.001). Baseline HAMD-17 and MADRS scores were highly correlated (r=0.817, p<0.001), as were the percent-change scores from baseline to post-treatment (r=0.948, p<0.001). MADRS was used as the primary clinical outcome for all FC analyses, and all results are equivalent for HAMD-17.

### Functional connectivity changes

We examined FC changes that occurred with SAINT. Paired t-tests (FDR-corrected) demonstrated significant decrease in FC between lsgACC and 3 of 4 DMN nodes (lsgACC-fDMN: t_(17)_=2.335, q =0.036; lsgACC-mDMN: t_(17)_=3.027, q=0.008; lsgACC-rDMN: t_(17)_=2.660, q=0.017). A similar trend was found for FC between lsgACC and lDMN (t_(17)_=2.106, q=0.06) (Figure 1).

**Figure 1.**
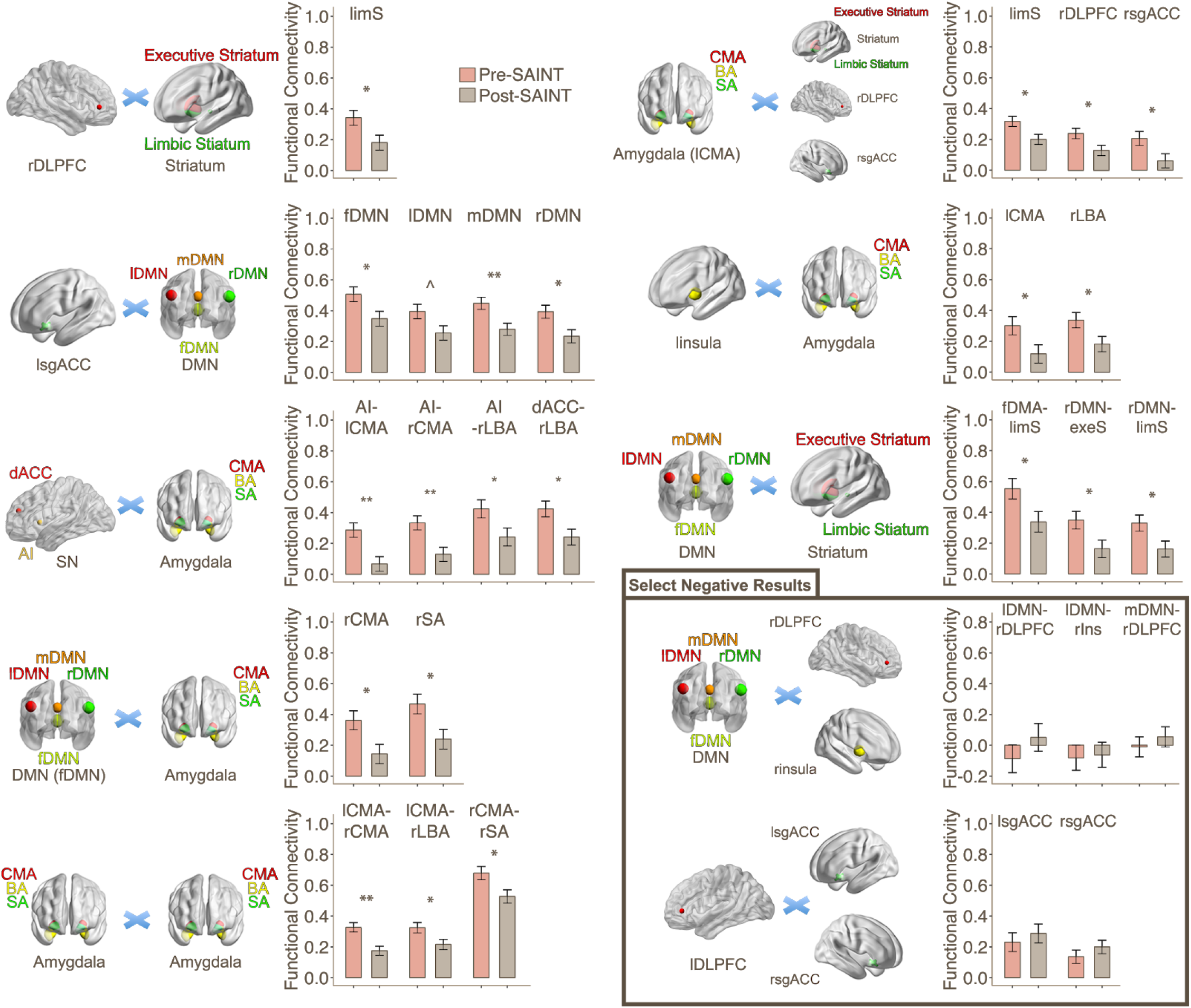
Statistically-significant changes in functional connectivity following SAINT in 18 participants with moderate-to-severe, treatment-resistant major depressive disorder. Bottom right box: Select non-statistically-significant results demonstrating observed negative FC with DMN and increase in FC between lDLPFC and sgACC. DLPFC: dorsolateral prefrontal cortex, sgACC: subgenual anterior cingulate cortex, DMN: default mode network, SN: salience network, dACC: dorsal anterior cingulate cortex, AI: anterior insula, CMA: centromedian amygdala, BLA: basolateral amygdala, SA: superficial amygdala. Preceding r: right, preceding l: left. Paired t-tests were corrected for multiple comparisons using FDR (q<0.05). Error bars indicate within-subject error. **q<0.01, *q<0.05, ^q=0.06

FC reductions were also observed between rDLPFC and amygdala (t_(17)_=2.303, q=0.036), and striatum (rDLPFC-limS: t_(17)_=2.354, q=0.032), between SN nodes and amygdala (AI-lCMA: t_(17)_=3.601, q=0.002; AI-rCMA: t_(17)_=3.167, q=0.006; dACC-rBLA: t_(17)_=2.515, q=0.023; AI-rBLA: t_(17)_=2.206, q=0.045), between DMN and amygdala (fDMN-rCMA: t_(17)_=2.491, q=0.024; fDMN-rSA: t_(17)_=2.520, q=0.023), between DMN and striatum (fDMN-limbic striatum: t_(17)_=2.281, q=0.039; rDMN-executive striatum: t_(17)_=2.303, q=0.037; rDMN-limbic striatum: t_(17)_=2.279, q=0.039), between left insula and amygdala (linsula-lCMA: t_(17)_=2.177, q=0.049, linsula-rBLA: t_(17)_=2.221, q=0.045), between amygdala and striatum (lCMA-limbic striatum: t_(17)_=2.515, q=0.023), and between amygdala sub-regions (lCMA-rCMA: t_(17)_=3.601, q=0.002; lCMA-rBLA: t_(17)_=2.276, q=0.024; rCMA-rSA: t_(17)_=2.481, q=0.040) (Figure 1). Select non-significant results are also included in Figure 1.

### Functional connectivity changes associated with clinical changes

We assessed the relationship between changes (percent change from baseline) in FC and depression symptoms using standard linear regression techniques adjusted for multiple comparisons (FDR).

Changes in FC between lDLPFC and other ROIs were significantly associated with changes in MADRS scores with better clinical outcomes correlated with greater decreases in FC between lDLPFC and lCMA (R^2^=0.210, t_(17)_=2.351; q=0.033) and between lDLPFC and linsula (R^2^=0.228, t_(17)_=2.452; q=0.026). Similarly, changes in FC between insula and amygdala were significantly associated with changes in MADRS scores with better clinical outcomes correlated with greater decreases in FC between rinsula and rBLA (R^2^=0.215, t_(17)_=2.377; q=0.031) (Figure 2).

**Figure 2.**
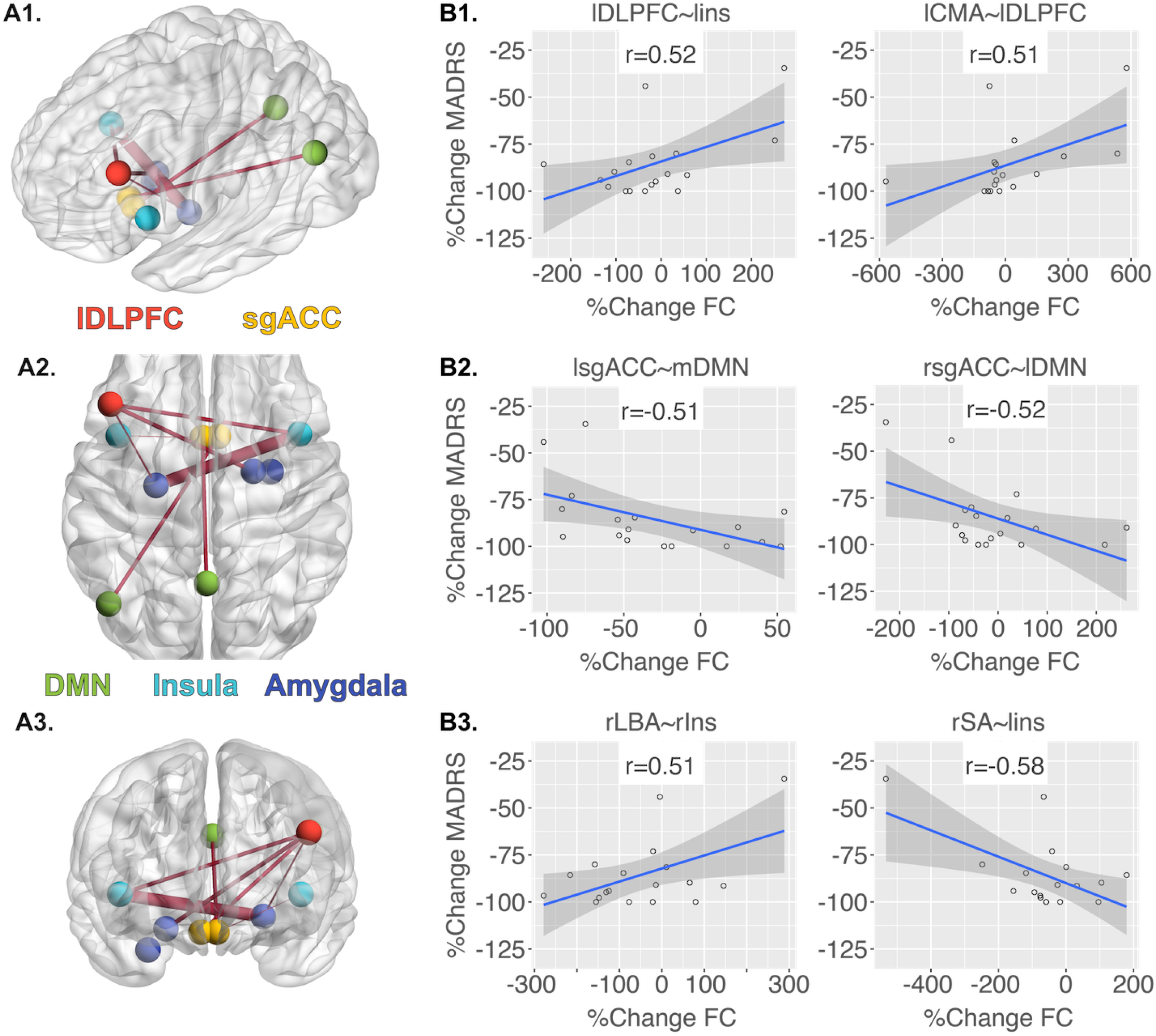
Statistically significant associations between changes in MADRS scores and FC changes following SAINT in 18 participants. **A**: Visualizations of the associations between changes in MADRS and changes in FC. The edges represent adjusted R^2^ values of the regression model - the greater the R^2^, the thicker the edge. Nodes are coded by color with arbitrary size. **A1**: sagittal image, **A2:** axial image, **A3:** coronal image. **B:** FC changes significantly associated with MADRS changes were observed between lDLPFC and insula, and amygdala subregions **(B1)**, between sgACC and DMN nodes **(B2)**, and between insula and amygdala subregions **(B3)**. FC: functional connectivity, MADRS: Montgomery Asberg Depression Rating Scale, DLPFC: dorsolateral prefrontal cortex, sgACC: subgenual anterior cingulate cortex, DMN: default mode network, CMA: centromedian amygdala, BLA: basolateral amygdala, SA: superficial amygdala. Preceding r: right, preceding l: left. The 95% confidence region is depicted by gray shading. All results were corrected for multiple comparisons using false discovery rate (q<0.05). %Δ = (*postTreatment – baseline*)/*baseline* * 100

In contrast, smaller magnitude reductions (and larger magnitude increases) in FC between sgACC and DMN nodes were associated with better clinical outcomes. Specifically, greater depression symptom improvements were correlated with increased FC between rsgACC and lDMN (R^2^=0.225, t_(17)_=-2.437, q=0.027) and FC between lsgACC and mDMN (R^2^=0.219, t_(17)_=-2.399, q=0.029). Although the overall group mean FC decreased with SAINT (see prior section), larger magnitude percent-reductions in MADRS scores were associated with smaller reductions and larger increases in FC between sgACC and DMN. The same relationship was found for FC between linsula and rSA (R^2^=0.291, t_(17)_=-2.827, q=0.012) (Figure 2).

Gender and age were not found to be confounders (all p’s>0.05), and all significant effects remained when including gender and age as covariates in multiple-regression models (all q’s<0.05).

### Baseline functional connectivity association with clinical changes

We assessed the association between baseline FC and changes in MADRS scores with a similar linear regression approach. Larger improvements in MADRS scores were correlated with lower baseline FC between lDMN and frontal cortex (rVLPFC-lDMN: R^2^=0.235, t_(17)_=2.496; q=0.024; rDLPFC-lDMN: R^2^=0.231, t_(17)_=2.470; q=0.026) and between lDMN and striatum (limbic striatum-lDMN: R^2^=0.312, t_(17)_=2.951; q=0.009; executive striatum-lDMN: R^2^=0.306, t_(17)_=2.917; q=0.010) (Figure 3).

**Figure 3.**
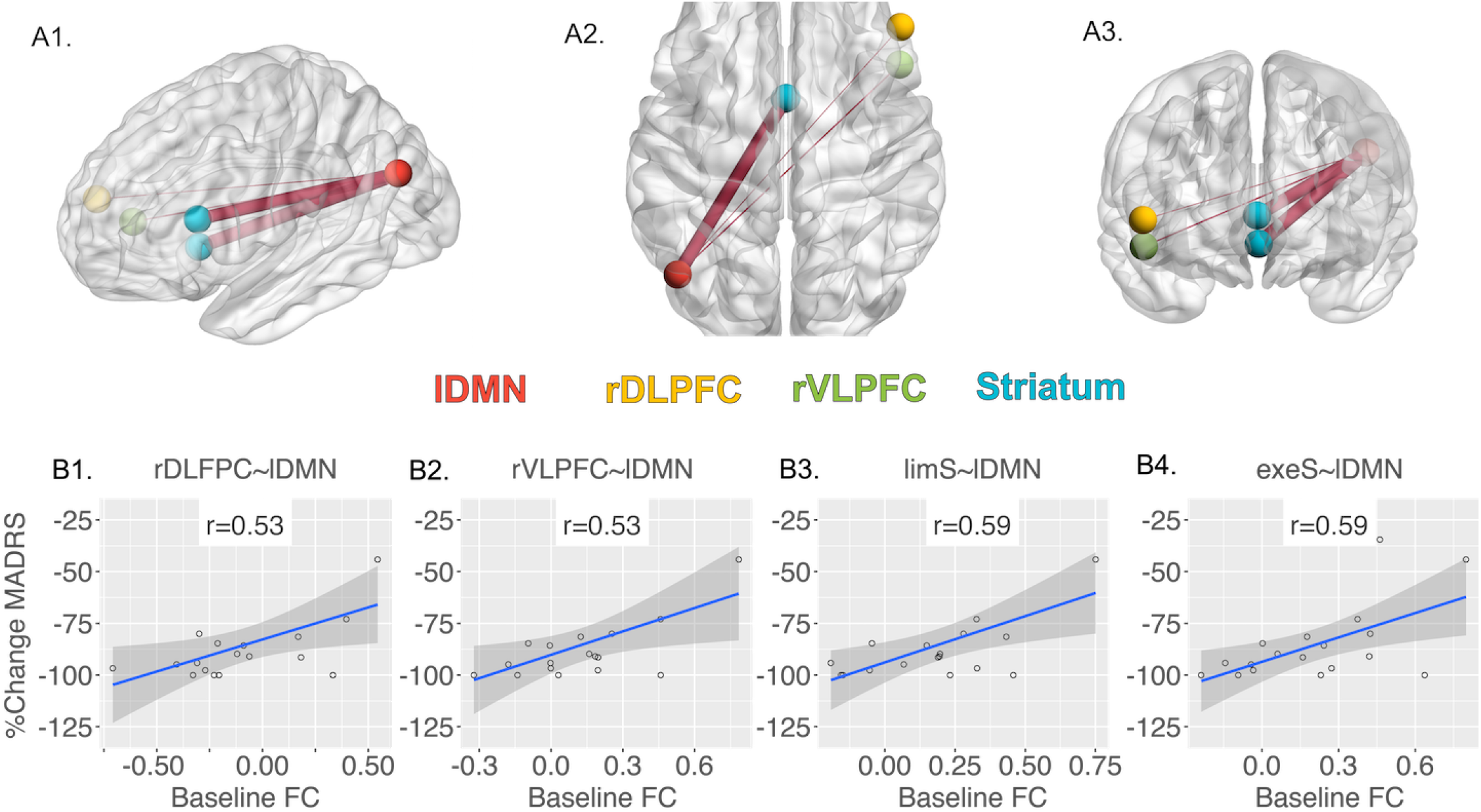
Baseline FC association with MADRS scores following SAINT. **A:** Visualizations of the associations between baseline FC and MADRS scores. The edges represent the adjusted R^2^ values of the regression model - the greater the R^2^, the thicker the edge. Nodes are coded by color with arbitrary size. **A1:** sagittal image, **A2:** axial image, **A3:** coronal image. **B:** Baseline FC between lDMN and frontal cortex **(B1, B2)**, and between lDMN and striatum subregions **(B3, B4)** were significantly associated with MADRS changes. FC: functional connectivity, MADRS: Montgomery Asberg Depression Rating Scale, DLPFC: dorsolateral prefrontal cortex, DMN: default mode network, VLPFC: ventrolateral prefrontal cortex, limS: limbic striatum, exeS: executive striatum. Preceding r: right, preceding l: left.The 95% confidence region is shaded gray. All results were corrected for multiple comparisons using false discovery rate (q<0.05). %Δ = (*postTreatment – baseline*)/*baseline* * 100

Gender and age were not found to be confounders (all p’s>0.05), and all significant effects remained when including gender and age as covariates in multiple-regression models (all q’s<0.05).

### Comprehensive results

Please see Supplementary Figure S1 for heat maps depicting all FC changes that were observed following SAINT. All pairwise comparisons for changes in FC following SAINT can be found in Supplementary Table 1. All linear regressions between change in FC and changes in MADRS are reported in Supplementary Table 2. All linear regressions between baseline FC and changes in MADRS are reported in Supplementary Table 3.

## Discussion

The development of SAINT has provided us with a unique opportunity to assess the FC changes that occur during rapid remission from moderate-to-severe, treatment-resistant MDD. The analysis we undertook in this initial report focused on brain regions associated with emotion regulation, and we found that FC significantly decreases between several of these regions following remission from MDD, with the magnitude of FC change often correlating with symptom improvement. Moreover, we found pre-treatment connectivity between several regions to be predictive of the magnitude of this improvement.

Notably, we observed a significant reduction in mean sgACC-DMN connectivity after SAINT, and this reduction was present between sgACC and 3 of 4 DMN nodes, with a nearly significant trend in the remaining node. In previous studies of depression, hyper-activity/connectivity within DMN has been considered as evidence of dysfunction of rumination and self-referential processing, and antidepressant treatment is hypothesized to normalize DMN activity and connectivity (24, 37, 65–67). Our results support this hypothesis and raise another important question: Does the magnitude of overall suppression of DMN FC track with antidepressant treatment efficacy? Prior studies have reported significant reductions in FC between sgACC and 1 to 2 nodes of DMN (24, 37, 65–67); whereas, we observed significant reduction in FC in nearly all DMN nodes. Bearing in mind that open-label SAINT was associated with remission in 90% of participants in 1-5 days (21), this question deserves careful consideration in future studies.

Moreover, this hypothesis builds on some of the earliest neuroimaging studies in the field of psychiatry, which reported increased sgACC activity in patients during depressive episodes (68). This has been hypothesized to be causally related to reductions in lDLPFC activity (69) and reinforced by observations that antidepressant treatments lead to reductions in sgACC activity (70–77). For review, see Hamani et al (78). Early investigators concomitantly reported observations of hypometabolism and hypoperfusion in the lDLPFC of patients with MDD (79–81). Later investigators demonstrated that lDLPFC activity is restored with treatment of MDD (70, 71, 82–87), and this early data prompted investigators to begin hypothesizing a major role of top-down dysfunction in the pathophysiology of depression (88, 89).

This led to the conception of depression as dysregulation in corticolimbic systems (89). In recent years, this concept has continued to evolve to include neural networks and their interactions. Several groups have reported evidence that depression is associated with increased FC between sgACC and DMN (35, 36, 53, 90–94), and others have shown that stimulation of lDLPFC potentiates upstream neural circuits to normalize sgACC-DMN connectivity (24, 37, 65–67). This does not seem to be limited to individuals with depression, as rTMS of lDLPFC also has been shown to improve mood and reduce connectivity between sgACC and DMN in never-depressed participants (95). Thus, we hypothesize that the significant reduction in FC in nearly all DMN nodes that we observed in SAINT may reflect the normalization of DMN hyper-connectivity associated depression and in part underlie its remission.

Interestingly, at the group level, FC between sgACC and DMN significantly decreased following SAINT in our cohort; however, individual changes in FC magnitude showed that participants with increased FC between sgACC and DMN actually had the greatest clinical improvement. One interpretation is that “normalization of FC between sgACC and DMN” is more complex than magnitude of FC change between 2 nodes. Alternately, disease heterogeneity may be related to changes in FC that are challenging to predict at the individual level with current approaches. Others have reported the opposite direction of association with the largest improvements in depression being associated with the largest reductions in FC between sgACC and mDMN (67); however, their patient cohort had comorbid post-traumatic stress disorder, which may respond differently to rTMS of the lDLPFC (96). Further, both Philip et al (67) and our current report found an overall reduction in mean FC between sgACC and DMN in participants with improved depressive symptoms. This possible discrepancy could also be due to the limited variability in post-treatment MADRS scores in our sample. Also, we do not know how FC changed throughout the treatment course.

Those who responded the earliest generally had larger reductions in MADRS scores; thus, there could be a staged, non-monotonic change in FC whereby FC initially decreases until symptoms remit and then increases with continued treatment. A similar non-monotonic time course has been reported for psilocybin, whereby acute administration decouples sgACC from DMN (97), but this connectivity increases from baseline in the days following treatment (18). Thus, additional studies will be crucial to determine the validity, reliability, and meaning of this association.

Our assessment of the association between baseline FC and clinical improvements in depression scores also revealed a central role of the DMN. Greater clinical improvements were associated with lower baseline FC between DMN and DLPFC, striatum, and ventrolateral prefrontal cortex. Although FC between DLPFC and DMN predicts efficacy of ECT (98), other studies with similar findings could not be located with extensive database searching. Past studies that have utilized FC to predict response have found that FC between stimulation site lDLPFC and striatum predicts TMS (23, 25), and several groups have reported that FC between stimulation site and sgACC that is more negative results in greater reductions in depressive symptoms with rTMS (28, 29, 59). In our current study, we optimized targeting by stimulating the portion of lDLPFC with maximum anticorrelation with sgACC; thus, it is possible that this may underlie the striking reductions in depression symptoms we observed. Finally, in contrast to antidepressant medications (30), intact baseline connectivity between DMN and ACC was not related to outcomes following SAINT. Considering that depressive disorders may be distinguished based on intrinsic connectivity (99), it is interesting to note that SAINT was largely effective regardless of intrinsic connectivity, which may indicate that its mechanism of action affects circuits shared between subtypes of depressive disorders.

In addition to reduced FC between sgACC and DMN, we observed several other notable changes in FC. Specifically, we found reductions in connectivity between DLPFC and amygdala, insula, and limbic striatum to be associated with clinical improvement in depression. For rDLPFC-left amygdala FC, in addition to finding reduced positive FC, we also found that greater clinical improvement was associated with larger decreases in FC. Several fMRI studies have demonstrated that individuals with MDD have altered frontal connectivity with amygdala (40–44). Depressed individuals may have increased positive connectivity between rDLPFC and left amygdala (100, 101), and individuals with depression have an impaired ability to regulate negative emotions (41). Considering that non-depressed individuals can better suppress amygdala activity (43, 44), our results here could represent a partial normalization of connectivity related to improved emotion regulation, i.e. baseline positive FC between rDLPFC-amygdala is pathological, a reduction in positive FC is improved, and negative FC is normalized; however, our experimental design did not include never-depressed participants, so this remains speculative. Our combined findings of changes in FC with sgACC and amygdala following DLPFC stimulation with SAINT is supported by recent work with interwoven TMS and fMRI that showed DLPFC stimulation engages these nodes (102).

Similarly, we found that greater clinical improvements were associated with larger decreases in FC between lDLPFC and insula. Insula has a well-known role in interoception (103, 104), and insula activity is correlated with abnormal interoception in MDD (105). Insula metabolism has been linked to predicting antidepressant treatment response, and has thus been proposed as a depression biomarker (106). Our results here are similar to other reports. Theta-burst rTMS stimulation of lDLPFC has been reported to reduce the magnitude of FC between lDLPFC and right insula (107) with baseline FC between these structures predicting clinical response (108). Interestingly, individuals with subthreshold depression have lower FC between DLPFC (laterality not specified) and left insula compared to non-depressed individuals (109). This apparent directional mismatch highlights the importance of determining the causality of mechanism using within-subject analyses prior to using these findings as biomarkers to optimize treatment efficacy.

We also observed reduced FC in rDLPFC-limbic striatum following SAINT. Increased rDLPFC-dorsal caudate connectivity has been reported to correlate with increased depression severity (110), and FC between rDLPFC and ventral rostral putamen has been shown to decrease following depression treatment (111). Similarly, Furman et al found that depressed individuals have greater FC between DLPFC and dorsal caudate, and this FC was significantly correlated with depression severity, but they did not specify laterality (45). This association has also been reported for lDLPFC-ventral striatum FC. Avissar et al reported that improvement in depression correlated with more reduction in lDLPFC-ventral striatum FC following treatment with rTMS of lDLPFC (23). Thus, reducing DLPFC-striatum FC may be an important route to inducing remission from depression. The combined results show consistent association between significant changes in DLPFC FC with limbic structures following SAINT, which supports the hypothesis that regulation of emotion plays a central role in MDD.

This hypothesis is further supported by our observation that following SAINT we found a significant decoupling of SN nodes, i.e. reduced FC between amygdala-insula, amygdala-dACC, amygdala-limbic striatum, and between amygdala subregions. Increased SN FC has been reported to be associated with decreased anticipated pleasure (112) and MDD (113). Further, other groups have reported that individuals with MDD have greater FC between insula and amygdala (105, 114) and amygdala and striatum (114). This dysfunction in SN FC may underlie the biased reactivity to negative emotions observed in individuals with MDD (113, 115).

Some results we observed are more challenging to interpret. For example, following SAINT, FC significantly decreased between fDMN and multiple subregions of right amygdala and between multiple DMN nodes and striatum. rTMS of the lDLPFC has previously been shown to reduce FC between DMN and striatum in non-depressed individuals (95). However, other reports suggest the opposite association for mDMN (posterior cingulate cortex and precuneus): Individuals with depression have reduced FC between amygdala and precuneus compared to non-depressed controls (116), and connectivity between amygdala and posterior cingulate cortex is greater in less depressed individuals (117). Our finding that FC between DMN and striatum decreased after SAINT is consistent with groups who have observed a similar change with antidepressant (118) and ECT treatment (119). Others have reported that FC between mDMN and caudate is reduced in depressed individuals (120), indicating this may be a compensatory mechanism; however, this is complicated by reports that reduced FC between DMN and ventral striatum is associated with impaired reward responsivity (121). Thus, a causal association between these changes in FC with depression is difficult to infer at this time.

Our observations should be considered in the context of the study’s weaknesses. Foremost, the open-label design without a sham or standard-of-care rTMS control group limits our ability to determine if FC changes were due to SAINT, remission from MDD, or the passage of time. Sham-controlled, double-blinded assessment of SAINT is currently underway to address this important question. A strength and weakness of our analysis is the low variability in treatment-associated changes in depressive symptoms. Ninety percent of individuals were in remission from MDD following SAINT, with only 2 participants not remitting in this cohort, making statistical comparisons between remitters and non-remitters non-informative. Additionally, our linear-regression assessments demonstrated floor-effects, likely eliminating the potential to observe possible associations between changes in FC and clinical response. The small sample size further limited our likelihood of observing inter-individual differences. On the other hand, the high proportion of remitters increases our confidence in mean changes in functional connectivities across the group. Follow-up controlled trials will be helpful in determining the reliability of the findings we observed here. A limitation of our neuroimaging methods is that our resolution was insufficient to explore the role of the hippocampus and its sub-regions. As the hippocampus has numerous reports describing an important role in depression (122), future studies of SAINT would benefit from assessment of FC related to this important structure. Finally, we included ROIs with multiple prior reports demonstrating a role in emotion regulation; however, we did not confirm that the changes in FC that we observed were associated with changes in emotion regulation beyond standardized depression scales.

Depression remains a major public health problem with current treatments taking weeks to months to induce remission, and there are millions of people who are not effectively treated with current methods. Development of improved treatments could be accelerated by a better understanding of the neural changes that are necessary and sufficient to achieve disease remission. Our recent development of SAINT, which produces rapid and high rates of remission from MDD, provides an opportunity to begin searching for these neural changes. Our initial analysis of brain regions associated with emotion regulation demonstrates MDD remission is associated with a significant reduction in FC between sgACC and almost all major nodes of the DMN, bolstering prior hypotheses of the central role this network plays in MDD. We also found that decoupling of SN nodes was correlated with depression improvement, highlighting the role of other networks as well. As our current study is based on an open-label trial of SAINT, controlled trials will be necessary to determine which observed changes in FC are unique to SAINT and disease remission in order to prioritize network-level targets for further treatment optimization.

## Materials and Methods

### Participants

Detailed information regarding the participants, SAINT protocol, experimental design, and behavioral data analysis can be found in our prior report (21). Briefly, participants treated with SAINT had a diagnosis of treatment-resistant MDD and complete MRI scans both pre- and post-SAINT. Diagnosis of MDD was confirmed with the Mini International Neuropsychiatric Interview (MINI). Participants were required to have a Hamilton Depression Rating Scale 17-item (HDRS-17) score higher than 20 and not have responded to at least 1 antidepressant medication (minimum trial duration of 6 weeks) to be eligible for the study. Participants had no history of a psychotic disorder, substance use disorder, major systemic illness, current mania (assessed using the Young Mania Rating Scale), lesional neurological disorder, or any contraindications to rTMS or MRI (such as implanted metal or history of seizures). All participants provided written informed consent, and the study was approved by the Stanford University Institutional Review Board and in accordance to the Declaration of Helsinki.

### Stanford Accelerated Intelligent Neuromodulation Therapy (SAINT)

Participants received a 5-day treatment course under the SAINT protocol. Stimulation location was chosen based on an algorithm that selected the area within the left dorsolateral prefrontal cortex (lDLPFC) with the most negative FC with sgACC. This is described in detail in our prior report (21). A MagVenture MagPro X100 (MagVenture A/S, Denmark) system was used to deliver sessions of intermittent theta-burst stimulation (iTBS): 60 cycles of 10 bursts of 3 pulses at 50Hz were delivered in 2 second trains (5Hz) with an 8 second inter-train interval. Stimulation sessions were delivered hourly. Ten sessions were applied per day (18,000 pulses/day) for 5 consecutive days (90,000 pulses in total). Stimulation was delivered at 90% resting motor threshold (rMT). A depth correction (64) was applied to the resting motor threshold to adjust for differences in the cortical depth of the individual’s functional target compared to the primary motor cortex in order to consistently achieve 90% rMT in the intended functional target, but stimulation was never delivered above 120% rMT for safety. A Localite Neuronavigation System (Localite GmbH, Sankt Augustin, Germany) was used to position the TMS coil over the individualized stimulation target. All participants in this report’s subset completed the full 5-day treatment course.

### Clinical assessment

Participants’ depressive symptoms were assessed using the Hamilton Depression Rating Scale-17 item (HAMD-17) (123) and Montgomery-Asberg Depression Rating Scale (MADRS) (Montgomery and Asberg, 1979). HAMD-17 and MADRS assessments were performed 1-4 days before SAINT (baseline) and 1-4 days after the 5-day SAINT treatment (post-treatment) in concordance with MRI scans.

### MRI image data acquisition and preprocessing

All participants were screened for MRI safety prior to any scanning procedures. MRI scans were acquired approximately 72hrs before SAINT (baseline) and approximately 72hrs after the 5-day SAINT treatment (post-treatment). Each participant underwent identical baseline and post-treatment MRI scans consisting of structural and resting-state functional MRI acquisitions. All MRI scans were acquired using a 3T GE Discovery MR750 scanner with a 32-channel head-neck imaging coil at the Center for Cognitive and Neurobiological Imaging at Stanford.

High-resolution structural images using GE’s “BRAVO” sequence (three-dimensional, T1-weighted) were acquired for the whole brain (FOV=256×256mm; matrix=256×256 voxel; slice thickness=0.9mm; TR =2530ms, TE = 2.98ms, flip angle=7°).

During the 8-min resting state scans, participants were instructed to keep their eyes open and their attention focused on a central fixation point, which consisted of a black screen with a white fixation cross. Participants were also instructed to let their minds wander freely and to avoid any repetitive thoughts such as counting. Whole-brain resting-state scans were collected with a 3X simultaneous multi-slice (i.e. multiband) acquisition echo planar imaging (EPI) sequence: TR = 2000ms, TE = 30ms, flip angle = 77°, slice acceleration factor=3, FOV=230×230mm, matrix=128×128 voxel, 1.8×1.8 mm^2^ in-plane resolution, 87 contiguous axial slices parallel to the anterior commissure - posterior commissure line, yielding >1.4M voxels every 2 seconds. Head motion of participants was effectively minimized with the use of memory foam and inflatable padding. Participant alertness during the resting state task was monitored using in-scanner video cameras.

The MRI data were preprocessed using FMRIPREPR version 1.2.3 (124) [RRID:SCR_016216]. Each T1-weighted volume was corrected for intensity non-uniformity with N4BiasFieldCorrectionv2.1.0 (125) and skull-stripped using antsBrainExtraction.shv2.1.0 (using the OASIS template). Spatial normalization to the ICBM 152 Nonlinear Asymmetrical template version 2009c (126) [RRID:SCR_008796] was performed through nonlinear registration with the antsRegistration tool of ANTs v2.1.0 (127) [RRID:SCR_004757], using brain-extracted versions of both T1-weighted volume and template. Brain tissue segmentation of cerebrospinal fluid, white matter, and gray matter, was performed on the brain-extracted T1-weighted volume using fast (128) (FSL v5.0.9, RRID:SCR_002823).

The resting state data were motion corrected and then transformed into standard space (MNI). Physiological noise regressors were extracted applying CompCor (129). Framewise displacement (130) was calculated using the implementation of Nipype. Unsteady state volumes detected by FMRIPREPR were censored out (range 3 ~ 9, mean=5.12, SD=1.15). Independent component analysis-based Automatic Removal Of Motion Artifacts was used to non-aggressively denoise the data (131). Next nuisance signal regression was applied with signals from white matter and cerebrospinal fluid. All the volumes with framewise displacement (FD) greater than 0.2mm were excluded; this resulted in exclusion of 1 participant, as greater than half of the dataset had FD>0.2mm. All the data were spatially smoothed (6-mm-full-width, half-maximal Gaussian kernel) and temporal bandpass filtered (0.1-0.01 Hz). Data were detrended using Nilearn (132).

### Regions of interest (ROI)

For this initial analysis, we restricted our ROIs to brain regions that have consistently been reported to play a role in MDD and emotion regulation: DLPFC, VLPFC, DMN, sgACC, SN, insula, amygdala and striatum.

In line with prior studies (26, 37, 78, 133, 134), the DLPFC (BA9, BA46), VLPFC (BA44, BA45, BA47), and the sgACC (BA25) were defined by Brodmann areas (BA), using the Talairach atlas (135). The 4 nodes of DMN were extracted from the Multi-Subject Dictionary Learning (MSDL) atlas. DMN nodes were labeled using MSDL nomenclature: fDMN (Frontal DMN: medial prefrontal cortex), mDMN (median DMN: posterior cingulate cortex and precuneus), lDMN (left DMN: left angular gyrus), rDMN (right DMN: right angular gyrus) (136). The nodes of the SN, including anterior insula (AI) and dorsal anterior cingulate cortex (dACC), and insula were also obtained from MSDL. The striatum and amygdala are large nuclei with heterogeneous functions (23, 137–141); hence, we constructed our ROIs of these structures by subregions. The executive and limbic subregions of striatum were defined using Oxford-GSK-Imanova Striatal Connectivity Atlas (striatum-con-label-thr25-3sub) (141). Amygdala and its subregions were extracted using the Julich histological atlas: centromedian amygdala (CMA), basolateral amygdala (BLA), and superficial amygdala (SA) (142).

### Data analysis

All statistical analyses were conducted with R (143). FC was calculated as the correlation (Pearson) of the activity pattern across time between ROIs. All Pearson correlation coefficients were then transformed into Fisher’s Z score for further analysis. Changes in mean MADRS, HAMD-17, and FC were assessed with paired t-tests. Changes in MADRS and FC were expressed as percent change from baseline: Delta=100*(post-treatment - baseline)/baseline; thus, negative values of MADRS changes indicate improvement in depression symptoms. Linear regression models were used to test the relationship between FC and depression scores with R^2^ values adjusted using the Wherry Formula (144). Statistical significance was corrected for multiple comparisons across all comparisons using false discovery rate (FDR, threshold q=0.05). This included correcting across 231 correlations between FC and MADRS changes, 231 correlations between baseline FC and MADRS changes, and 231 paired t-tests.

## Author Contributions

B.S.B and X.X. designed experiments, analyzed data, interpreted data analyses, and prepared the manuscript. E.J.C., C.T., K.H.S., and D.D. collected data. J.H.B., A.S., and C.K. assisted in interpreting data analyses. K.D.S co-designed and executed the algorithm used to define the optimal stimulation targets. N.R.W invented the stimulation methodology, co-designed the algorithm used to define the optimal stimulation targets, designed the trial, acted as study MD, and mentored the writing of the manuscript

## Trial registration

ClinicalTrials.gov NCT03240692

## Abbreviations

AI: anterior insula
BLA: basolateral amygdala
CMA: centromedian amygdala
dACC: dorsal anterior cingulate cortex
DLPFC: dorsolateral prefrontal cortex
DMN: default mode network
exeS: executive striatum
FC: functional connectivity
HAMD: Hamilton Depression Rating Scale
Ins: insula
limS: limbic striatum
MADRS: Montgomery Asberg Depression Rating Scale
MDD: major depressive disorder
SA: superficial amygdala
sgACC: subgenual anterior cingulate cortex
SN: salience network
VLPFC: ventrolateral prefrontal cortex.
Preceding r: right
preceding l: left.

## Acknowledgements

The authors kindly thank the following for supporting this study: Charles R. Schwab, The Marshall and Dee Ann Payne Fund, the Lehman Family, Neuromodulation Research Fund, Still Charitable Fund, Avy L. and Robert L. Miller Foundation, Stanford Psychiatry Chairman’s Small Grant, Stanford CNI Innovation Award, NIH T32 035165, NIH UL1 TR001085, Stanford Medical Scholars Research Scholarship, NARSAD Young Investigator Award, the Gordie Brookstone Fund, and the Stanford Department of Psychiatry & Behavioral Sciences.

## Figures

**Figure S1.**
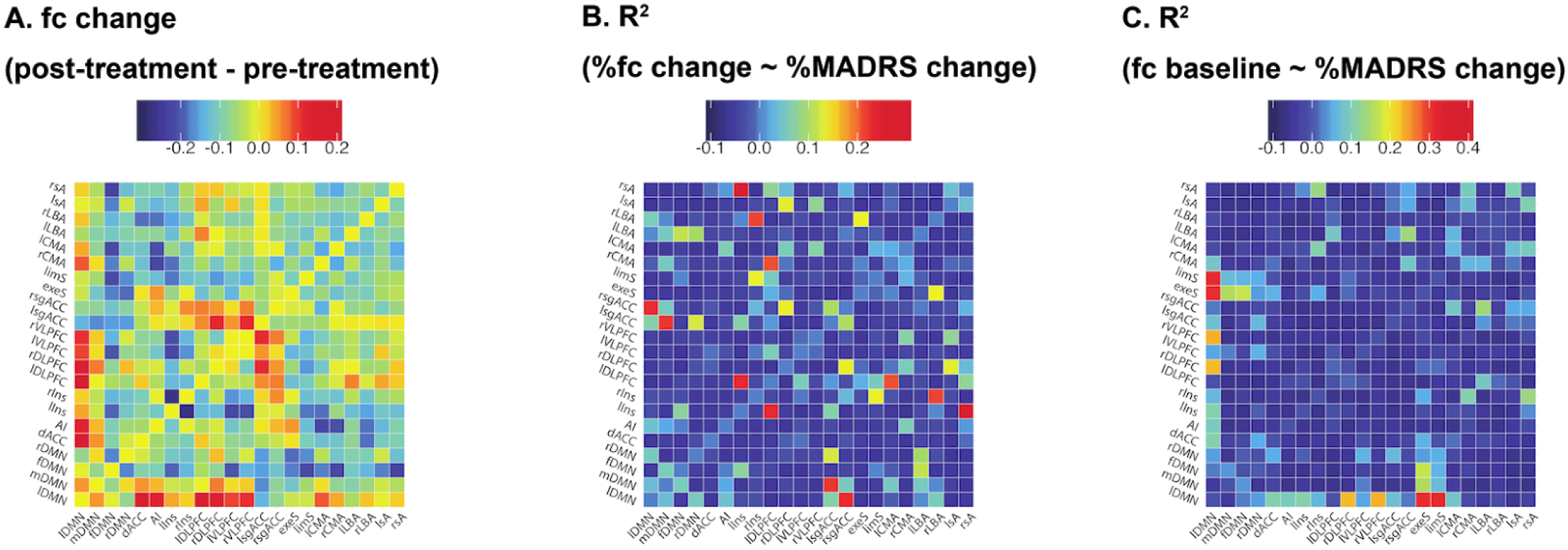
**A:** Functional connectivity changes (Δ= *postTreatment – baseline*) between each pair of all 22 ROIs. **B**: Adjusted R^2^ for the linear models relating functional connectivity changes to clinical score changes for each pair of all 22 ROIs. **C:** Adjusted R^2^ for the linear models relating baseline functional connectivity to clinical score changes for each pair of all 22 ROIs. %Δ = (*postTreatment – baseline*)/*baseline* * 100

**Table 1:**
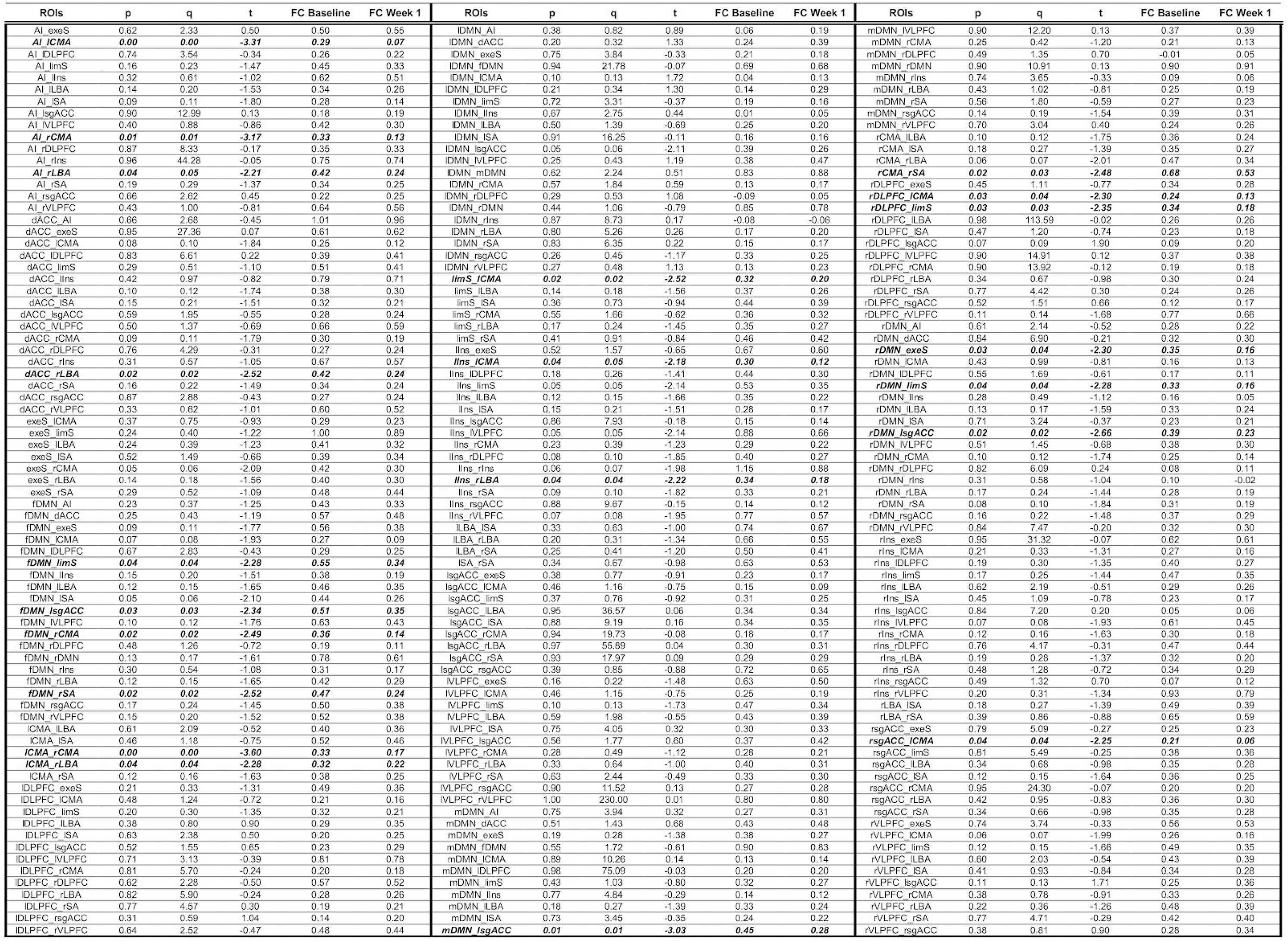
Mean FC values and paired t-test results for all ROIs.

**Table 2:**
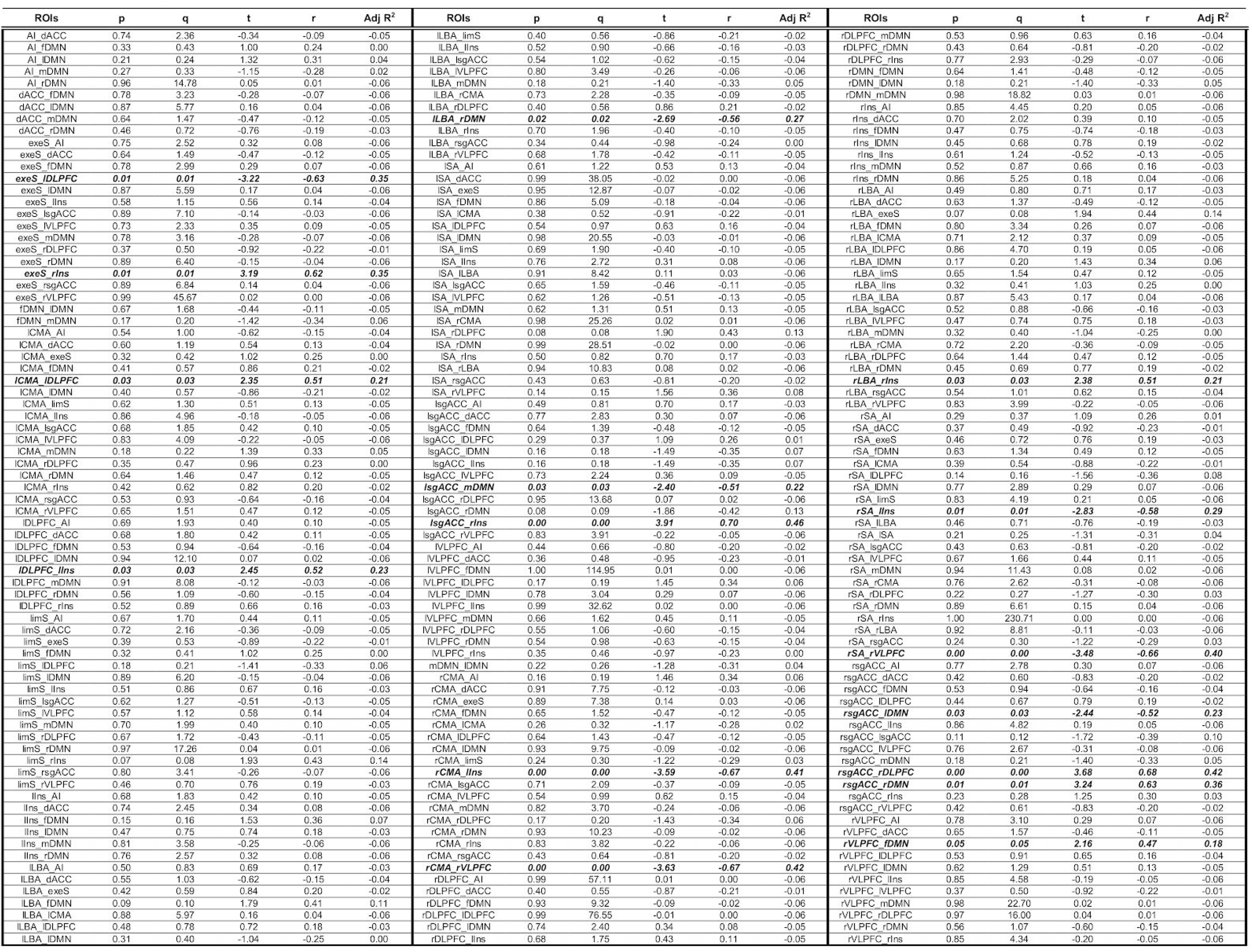
Linear regressions for %Δ FC vs. %ΔMADRS for all ROIs.

**Table 3:**
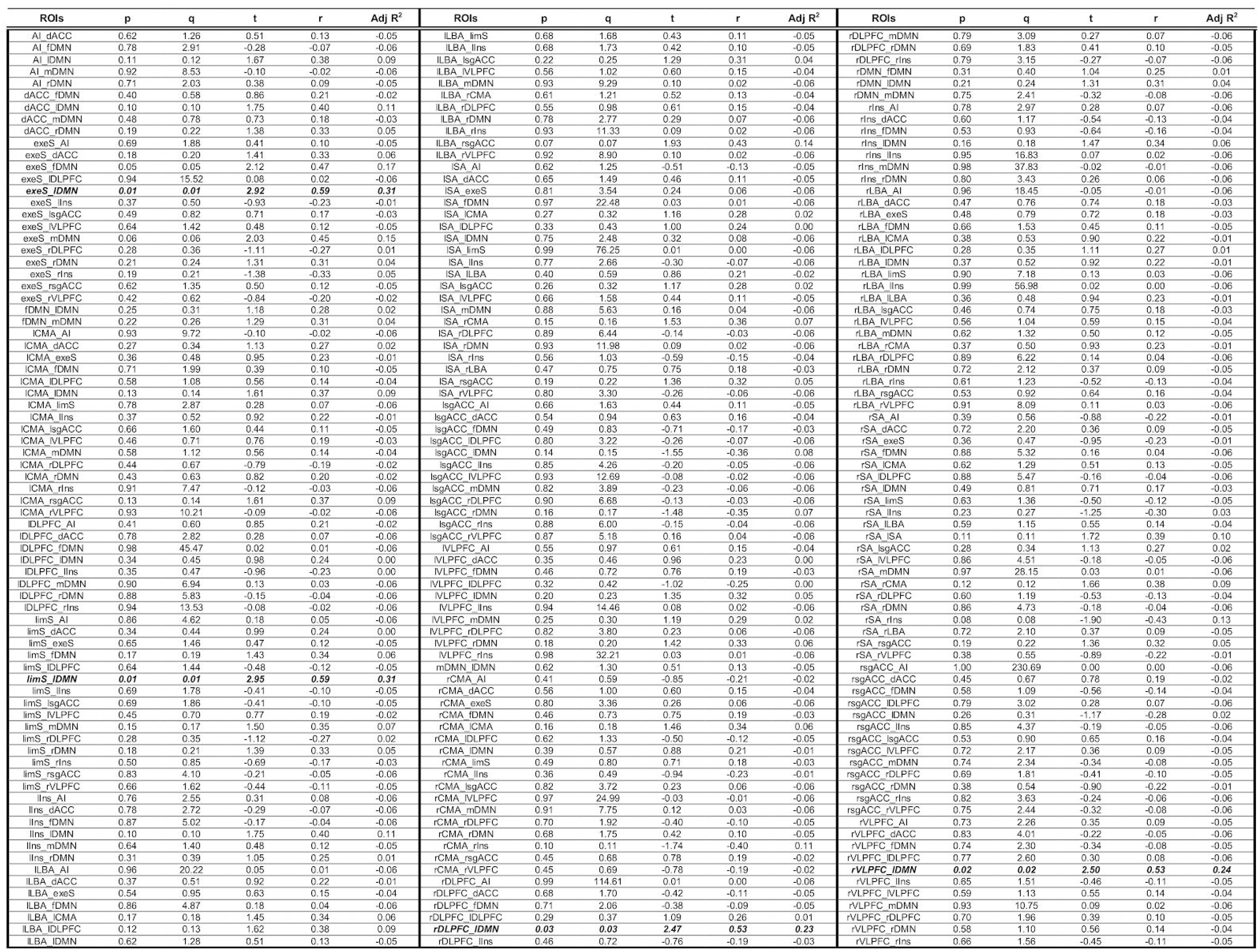
Linear regressions for baseline FC vs. %ΔMADRS for all ROIs.

